# Distinct cortical networks subserve spatio-temporal sampling in vision through different oscillatory rhythms

**DOI:** 10.1101/2022.05.09.491140

**Authors:** Luca Ronconi, Elio Balestrieri, Daniel Baldauf, David Melcher

## Abstract

Although visual input arrives continuously, sensory information is segmented into discrete events. Here, the neural correlates of spatiotemporal binding in male/female human subjects were investigated with MEG using two tasks where separate flashes were presented on each trial but were perceived, in a bi-stable way, as either a single, or two separate, events. The first task (two-flash fusion: TFF) involved judging one versus two flashes while in the second task (apparent motion: AM) participants judged coherent motion versus two stationary flashes. Results indicate two different functional networks underlying two unique aspects of visual temporal binding. In the TFF task, involving an integration window of ≈50 ms, evoked responses differed as a function of perceptual interpretation by ≈25 ms after stimuli presentation. Multivariate decoding of subjective perception based on prestimulus oscillatory phase was significant for alpha-band activity in the right medial temporal (MT) area, with the strength of pre-stimulus connectivity between early visual areas and MT being predictive of performance. In contrast, the longer integration window (≈130 ms) for AM showed evoked field differences only ≈250 ms after stimuli onset. Phase decoding of the perceptual outcome in the AM task was strongest for theta-band activity, localized to a right intra-parietal sulcus (IPS) source. Pre-stimulus connectivity between MT and IPS seeds in the theta band best predicted perceptual outcome. Overall, these results show a strong relationship between specific spatiotemporal binding windows and specific oscillations, linked to the information flow between different areas of the “where” and “when” visual processing pathways.

**Significance Statement:** Multiple neural rhythms seem relevant for sampling visual information across space and time, but the cortical networks underlying these fundamental computational principles of the visual system remain unexplored. We filled this gap by employing source-level multivariate decoding and connectivity analyses of magnetoencephalographic data recorded during an integration/segregation task of temporal and spatio-temporal events. We identified a first and faster network involving early visual areas (V2 to MT/V5) that determines the basic temporal resolution of visual perception at the speed of the alpha rhythm, and a second slower network involving parietal regions (IPS) that had a key role in the integration of more complex spatiotemporal events at a theta speed. These findings elucidate the neural mechanisms that transfer sensory information into temporal sequences.

## Introduction

Many aspects of our lives, including motion processing, speech recognition, reading, sound localization and visuo-motor coordination requires temporal or spatio-temporal integration and segregation of sensory information in the subsecond scale. This fundamental process represents a core mechanism of perception, allowing change in the flow of sensory input to be consciously represented without any experience of discontinuity (White, 2018).

After seminal neurophysiological investigations proposing that perception depends on the rhythmic sampling of sensory information (Bishop, 1932; Lansing, 1957; Harter, 1967), the neural correlates of spatiotemporal integration/segregation have been linked to ongoing neural oscillations using neurophysiological techniques in humans (Varela et al., 1981; Pöppel, 1997). The main hypothesis proposes that the alpha rhythm (8-12 Hz) defines a neural computation cycle within which integration of visual inputs occur (VanRullen, 2016). This idea is supported by studies in non-human primates showing that spikes in sensory areas are more likely to occur at a specific phase of the local field potential oscillations (such as the peak or trough) compared to opposing phases (Haegens et al., 2011). However, we also know that different sensory modalities have different preferential rhythms for organizing sensory evidence over time. For example, it is well established that visual information is sampled in a much narrower frequency band than auditory information due to the different neural architecture of peripheral and central information transmission (Morillon and Schroeder, 2015; White, 2018). It is reasonable to hypothesize that even within a single sensory modality, the sampling rhythm may vary according to the complexity of the temporal integration/segregation to perform. Perceiving temporal variation related to complex visual objects, such as words or faces, would require a more complex and extensive brain network as opposed to perceiving changes in a simple stimulus, such as an oriented line or grating (e.g., Baldauf and Desimone 2014; de Vries and Baldauf 2019). Similarly, integrating information about a dynamic event across space and time would involve a more complex network, and potentially a longer computation cycle, compared to a stationary stimulus.

Recent evidence has brought support in favor of the idea that within the visual modality there are multiple preferential neural rhythms for the sampling of sensory information across space and time. Specifically, the perceptual sampling of stimuli that alternate in close temporal proximity and in the same spatial position has been linked with alpha band activity (Cecere et al., 2015; Samaha and Postle, 2015; Gulbinaite et al., 2017), whereas stimuli separated by larger temporal intervals that also require sampling across space appear to involve slower frequencies within the theta band (Ronconi et al., 2017). This idea is in line with a theoretical framework of rhythmic perception that suggests that the frequency of ongoing neural oscillations determines the resolution of rhythmic sampling, in a way that higher oscillatory frequencies imply shorter temporal integration windows (Samaha and Postle, 2015; Ronconi and Melcher, 2017; Ronconi et al., 2018; Wutz et al., 2018).

However, a characterization of the networks supporting these different sampling mechanisms, which co-exist to determine our perception of the continuous sensory flow, is presently lacking. Specifically, there is a need for a precise mapping between the core spatiotemporal sampling mechanisms of human perception and the related rhythm-based cortical network dynamics. In the present study, we aimed to fill this gap by investigating the neural correlates of spatiotemporal sampling in humans using magnetoencephalography (MEG). In the same blocks of trials, participants performed two perceptual discriminations, a two-flash fusion and an apparent motion task, measuring temporal and spatiotemporal integration/segregation mechanisms, respectively. In both tasks, two separate flashes were physically presented on each trial, but participants perceived them in a bi-stable way. In the two-flash fusion condition, temporal integration would lead to the conscious report of a single stimulus as opposed to two discrete flashes, whereas in the apparent motion condition spatio-temporal integration would lead to a conscious report of single moving object as opposed to two discrete flashes in different spatial position.

## Methods

The main steps involved in the present study – and described in detail below – developed as follows: first, we mapped the cortical regions that differentiated integration versus segregation in the two different task conditions, by analyzing MEG activity evoked after the stimulus onset as a function of the subjective perceptual interpretation of the same bi-stable stimuli. Second, we trained a multivariate classifier to decode the perceptual outcome from the phase of pre-stimulus oscillatory activity within these networks. Finally, we used the resulting information to characterize the network-level interactions in terms of functional connectivity.

### Participants

30 participants (20 females), aged 18-35, took part in the study as paid volunteers. No participants reported history of neurological disease or epilepsy. All of them reported normal or corrected-to-normal vision and hearing and gave informed written consent. Three subjects were removed for the subsequent analyses, one because of excessive MEG artifacts, and two because they perceived apparent motion in 90% of trials or more, thus their perception could not be considered bistable. The experimental protocol was approved by the Ethics committee of the Center for Mind/Brain Science at University of Trento and conformed to the principles of the Declaration of Helsinki of 2013.

### Apparatus, stimuli and task procedure

The display system used for presentation of visual stimuli within the magnetically shielded room was a DLP projector (PROPixx, VPixx Technologies Inc., Saint-Bruno, QC, Canada) running at a refresh rate of 100 Hz, aimed at a translucent back-projection screen (projected screen size 510mm x 380mm) located in a dimly lit, magnetically shielded chamber at a viewing distance of 100 cm. The stimulus presentation methodology follows the one previously used in Ronconi et al. (2017) and depicted in Figure 1A.

**Figure 1:**
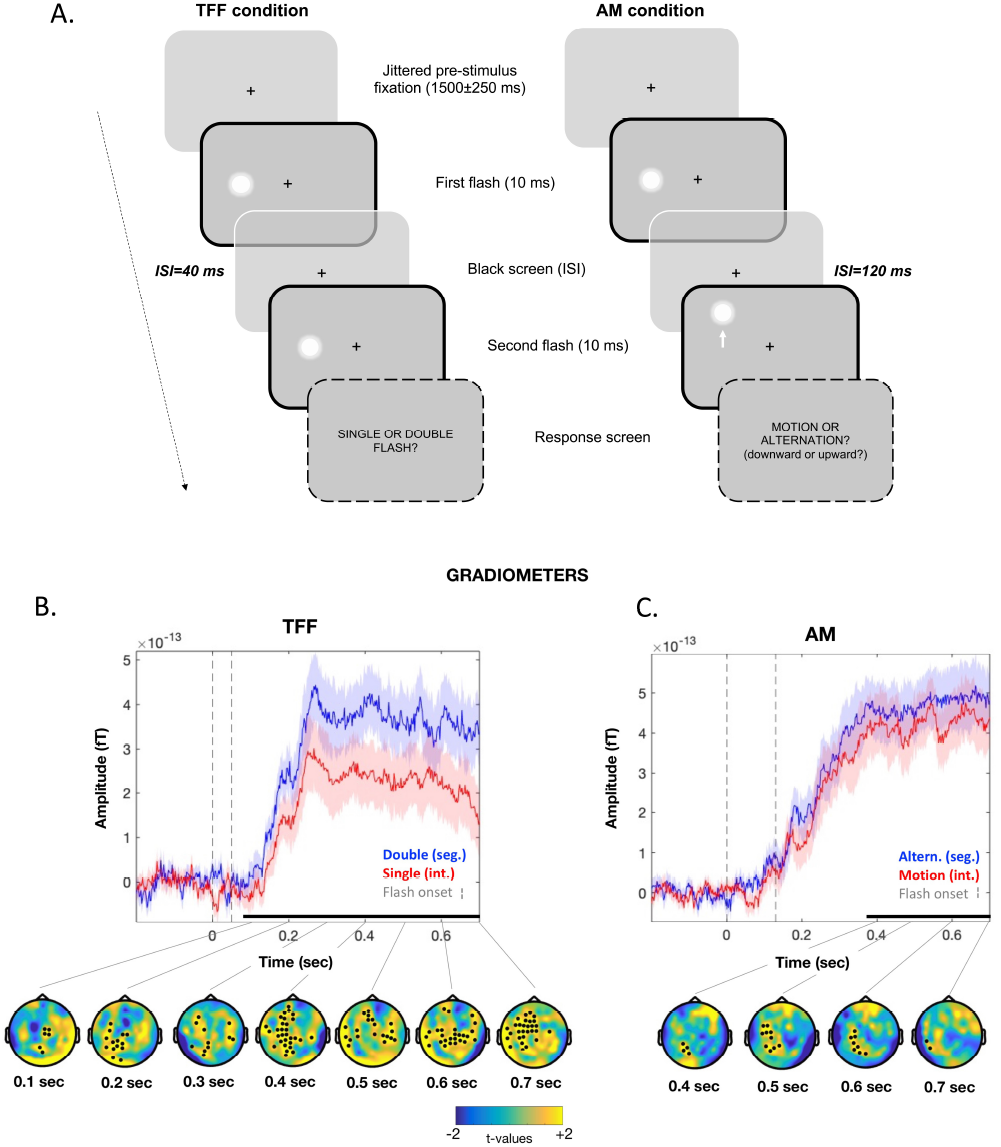
Task design and event-related fields (ERFs) results. (A) Schematic representation of the task procedure, where suprathreshold visual stimuli (flashes) were presented on the left or right hemifield either in a single (two-flash fusion or TFF condition) or in a different spatial position (apparent motion or AM condition), and participants were asked to judge if they perceive a single or moving stimulus or two distinct flashes. ISI=inter-stimulus interval. (B) (C) MEG gradiometers (average of all sensors) activity for the TFF (B) and the AM (C) conditions, differentiated as function of subjective perception (segregation vs. integration). Time points where significant cluster-corrected differences were found are highlighted with a black horizontal line above the time axis. Topographical maps at the bottom of each plot represent the time course of the segregation–integration difference, with black dots indicating significant clusters of channels.

Each trial began with a fixation point for a variable presentation time (ranging from 1350 to 1750 ms) and both target flashes had a duration of 10 ms (one refresh rate). In the two flash fusion (TFF) trials, the two target flashes appeared in the same position, aligned to the horizontal axis (left or right hemi-field, randomized across trials), with an eccentricity of 6 deg from the fixation. They were always separated by an inter-stimulus interval (ISI) of 40 ms (four refresh cycles).

In the apparent motion (AM) trials, the first of the two target flashes were again displayed at 6 deg of eccentricity aligned to the horizontal axis (left or right hemifield, randomized across trials). The second target flash appeared after an ISI of 120 ms (12 refresh rate) above or below the position of the first flash (at a distance of 4 deg) at the same eccentricity and in the same hemifield.

A blank screen of 1500 ms followed the target presentation, and anticipated the appearance of a response screen, in which participants had to report if they perceived one or two flashes for TFF trials, or if they perceived motion or alternation (and in which direction: upward or downward) for AM trials. No time constraints were imposed, and we stressed that only an accurate perception was important for the task and that reactions times were not relevant. After a response was entered, the subsequent trial started after an inter-trial interval of 1000 ms.

Each participant completed 10 MEG recording blocks of 8 minutes each, with an average number of trials completed equal to 751 (min. – max. range: 551-854). The different types of trials were randomly intermixed. An additional 5% of “catch” trials with longer ISI were presented for both trial types (100 ms for the TFF task and 200 ms for the AM task), with the aim of presenting clearly distinguishable targets that would reinforce bistable perception during the standard trials. Participants were unaware of the fact that bistable trials were all identical.

### MEG Data Acquisition

Participants’ whole-head MEG activity was recorded in a magnetically shielded room using a Neuromag 306 (Elekta) system with 102 magnetometers and 204 planar gradiometers, with a sampling rate of 1000 Hz. The system consisted of 102 sensors containing a triplet of one magnetometer and two gradiometers. To measure the head position while the participants’ head was within the MEG helmet, for each subject a specific headframe coordinates set was defined before the experiment, using pre-defined cardinal points of the head (i.e. nasion and left and right pre-auricular points), as well as the location of five head-position indicator (HPI) coils and a minimum of 200 other head-shape samples that were digitized for motion tracking using a Polhemus FASTRAK 3D digitizer (Fastrak Polhemus, Inc., Colchester, VA, USA). The subject’s head position relative to the MEG sensors was estimated before each MEG recording block (see Procedures) by activating the HPI coils to ensure that no major movements occurred during the data acquisition period.

### MEG Data analysis

Raw data were initially processed using MaxFilter 2.0 (Elekta Neuromag ^®^), which allows external sources of noise to be separated from head-generated signals using a spatio-temporal variant of signal space separation (tSSS) (Taulu and Kajola, 2005; Taulu et al., 2005). Before that, data were visually inspected and noisy channels were excluded from the tSSS filtering and replaced by interpolation. Movement compensation was applied and each run was aligned to an average head position.

After obtaining the Maxfiltered data, the subsequent data-analysis steps were performed in Matlab with the following freeware software packages: Fieldtrip for preprocessing, event-related fields and time-frequency analyses (Oostenveld et al., 2011), Brainstorm for cortical sources reconstruction (Tadel et al., 2011) and CoSMoMVPA for multivariate pattern analyses (Oosterhof et al., 2016).

Continuous MEG recordings were downsampled to 500 Hz and epoched from −1.5 s before to 1 s after the onset of the first stimulus. MEG epochs contaminated by artifacts were visually identified and manually rejected (an average of M = 21.05%, SD = 7.27% of trials for each participant were discarded after the artifact rejection procedure).

### Event-related fields (ERFs)

ERFs were calculated from artifacts-free epochs as the average in amplitude across trials, after combining data from planar gradient pairs using vector addition. ERFs were baseline corrected using an interval of −200 to 0 ms before the first stimulus onset. Statistical analyses between segregation and integration in the two tasks conditions were entirely data-driven; thus, we decided to perform permutations statistics (N=10000) and to apply a cluster-based correction for multiple comparison considering both time (all time points after stimulus onset) and sensor space (204 gradiometers) as dimensions to correct for, using a family-wise alpha level of .05.

Temporal windows where significant cluster-corrected differences emerged in the post-stimulus ERFs analyses where used to temporally constrain the identification of region of interest (ROIs) at the cortical source level, as described in the next paragraph.

### Cortical source reconstruction and ROI definition

The entire source reconstruction process followed the most recent guidelines for cortical sources reconstruction M/EEG data and related statistical analyses (Tadel et al., 2019). Structural magnetic resonance images (MRIs) were available for all participants (except one) and were all preprocessed with FreeSurfer (Fischl, 2012). For the only participant for which MRI was not available, we used the default cortical anatomy of the Montreal Neurological Institute (MNI).

We co-registered the brain surfaces from their individual segmented MRIs (Nolte, 2003) with an overlapping sphere head model. Empty-room recordings (2 minutes) collected the same day as the subject’s recordings were pre-processed following the same steps as participants’ data, and used to calculate the noise covariance matrix.

Forward modelling of electromagnetic fields was computed through the overlapping-spheres method (Huang et al., 1999). The estimation of distributed source amplitudes (inverse modelling) was computed using a weighted minimum-norm inverse kernel (wMNE) (Hämäläinen and Ilmoniemi, 1994). A z-score normalization was applied to each cortical source trace with respect to the prestimulus period (−200, 0 ms): this standardization replaces the raw source amplitude (pAm) value with new values that are suitable for hypothesis testing and, moreover, reduces the influence of inter-individual fluctuations in neural current intensity that is due to irrelevant anatomical or physiological differences (Tadel et al., 2019). Absolute values were used to compute the contrast measure between conditions regardless of the current’s polarity.

After obtaining the individual cortical maps of source activity for each individual, cortical sources were normalized onto a standard MNI brain (Montreal, Canada; http://www.bic.mni.mcgill.ca/brainweb). Surface smoothing was applied using a circularly symmetric gaussian kernel with a full width half maximum (FWHM) size of 5 mm. Such further steps improve the possibility to detect differential activity in a specific cortical region at the group level by reducing noise and inter-individual variability.

Finally, source data were averaged over the time points of interest that emerged from the ERF analyses and compared between the different subjective perceptual outcomes (segregation vs. integration), separately for both tasks. The resulting anatomical structures that were differentially activated as a function of subjective perception were labeled according to both the Desikan-Killiany and Brodmann atlases (see Table 1) and were used as ROIs for the MVPA of time-frequency data and functional connectivity (phase coherence) analyses, both described in the next paragraphs.

**Table 1:** Anatomical structures differentially activated as a function of subjective perception in the two task condition are listed together with their MNI coordinate (point of maximum difference); these cortical areas were labeled according to both the Desikan-Killiany and Brodmann atlases and were used as regions of interest (ROIs) for the MVPA of time-frequency data in the pre-stimulus interval and for functional connectivity analyses.

| Task | ROIs label | MNI coord | Cortical location (AAL) |
| --- | --- | --- | --- |
| TFF | V2.L | -3, -95, -24 | Cuneus.L |
|  | V2.R | 35, -86, -19 | Occipital_Inf.R |
|  | MT.R | 54, -70, 4 | Temporal_Mid.R |
|  | IFG.R | 53, 27, 8 | Frontal_Inf.Tri.R |
|  | MFG.R | 25, 58, 16 | Frontal_Sup.R |
|  | MFG.L | -37, 54, 26 | Frontal_Mid.L |
|  | IFG.L | -51, 19, -1 | Frontal_Inf.Tri.L |
|  | STG.L | -67, -12, 4 | Temp_Sup.L |
| AM | TPJ | 55, -47, 18 | Temporal_Sup.R |
|  | IPS/SupPariet.R | 36, -63, 42 | Angular.R |
|  | MTG.R | 47, -21, -8 | Temporal_Mid.R |
|  | Insula.R | 45, 1, 3 | Insula.R |
|  | SupFront.R | 27, -9, 69 | Frontal_Sup.R |
|  | SupFront.L | -18, -8, 78 | Frontal_Sup.L |
|  | IFG.R | 55, 18, -2 | Frontal_Inf.Oper.R |
|  | Insula.L | -43, -1, -4 | Insula.L |
|  | STG.L | -51, 18, -11 | Temporal_Pole_Sup.L |
|  | TempInf.R | 45, -47, -17 | Temporal_Inf.R |
|  | V1.L | -9, -95, 1 | Calcarine.L |

### Time-frequency decomposition and ROI-based single-trial phase decoding

Artifact-free epochs were transformed into time-frequency domain using a complex Morlet wavelet with varying number of cycles (3 at the lowest frequency and 10 at the highest) to obtain time-frequency (complex number) representation in 68 frequency bins from 3 to 30 Hz and 250 time points covering the entire epoch length relative to the stimulus onset.

Following a similar method used in our previous study (Ronconi et al., 2017), we used for each participant a searchlight with a cross-validated Naïve Bayes phase classifier to classify whether and at which frequencies the pre-stimulus phase of ongoing ROI activity could predict subjective perception. For the cross-validation, a split-half method was used on single trial source activity estimated for each ROI: 50% of the trials were selected pseudo-randomly for training the classifier, and the remaining half were used for testing. We performed this operation twice, training on one half and testing on the other half, and vice versa. Classification accuracy was computed as the number of correctly predicted condition labels divided by the total number of predictions. In all cases the train and test set were both balanced across the two conditions (integration or segregation). In other words, the number of trials in each condition was the same; where necessary (a few) trials were dropped using sub-sampling from the train or test set to ensure balance.

For the classifier, we used a custom re-implementation of some of the functionality present in the Circular Statistics Toolbox (Berens, 2009). We used a novel multivariate phase classification approach which was previously published in Ronconi et al. (2017). The input of the classifier was phase data from a set of trials with two conditions for a set of *k* features (combination of time points and frequencies). For each condition label c (indicating integration or segregation) and feature *i* in the training set, the average phase *θ_c,i_*, and concentration parameter *k_c,i_* was computed. For each trial in the test set, the probability *p_i,c_* that it belonged to class c according to feature *i* was computed using the von Mises circular probability density function (as implemented in circ_vmpdf.m in the circstat toolbox). Since our classification approach was Naïve Bayes (assuming independence across features), the combined class probability *P_c_* that a trial belonged to condition c was computed as *P_c_* = *p*_1,*c*_ * *p*_2,*c*_ *… *p_k,c_* integrating the information across the k features. The predicted condition label was set to the one with the highest probability. For improved accuracy when using very small probability values, our implementation took the logarithm of the probabilities and summed them. Since we used balanced trial counts for the two conditions there was no need to assume different prior probabilities accounting for class frequency.

For the temporal-frequency searchlight used in each ROI, each searchlight was based on radii of 4 time points and 8 frequencies. For a given ‘center’ feature (combination time point and frequency), features within a distance of 4 time points and 8 frequencies were selected and used for cross-validated classification as described earlier. The classification accuracy was then assigned to the center feature. This process was repeated for each feature, resulting in a classification accuracy map for all time points and frequencies within each ROI.

### Functional connectivity

Connectivity analysis was performed between pairs of ROIs defined at the cortical source level, using as hubs the specific ROIs where perception could be successfully decoded. Estimating functional connectivity at the source level has the advantage of reducing the effect of electromagnetic field spread and preventing spurious (non-independent) source-leakage effects, such as linear mixing or cross-talk between time series (Schoffelen and Gross, 2009). Specifically, we hypothesized that stronger connectivity states around the stimulus onset would lead to better communication between lower-order and higher-order visual regions, in agreement with recent findings (Rassi et al., 2019), thus promoting a more accurate representation (i.e. segregation) of visual stimuli.

To estimate the coupling between pairs of ROIs, we employed the magnitude squared coherence, a widely used measure of phase-dependent connectivity (Schoffelen and Gross, 2009), calculated in a pre-stimulus time period extending 1 second before the first stimulus onset. As before, we used the same number of trials to estimate connectivity in the two conditions (integration or segregation), by subsampling the condition with more trials.

We focused our analysis in the frequency bands which emerged as significant predictors of subjective perception in the pre-stimulus phase decoding analyses. Based on our previous study (Ronconi et al., 2017) we expected these frequencies to be in the theta and alpha band. Given that the frequency of alpha could play a role in determining integration vs. segregation of visual stimuli (Samaha and Postle, 2015; Ronconi et al., 2018; Wutz et al., 2018), the whole alpha band was split into lower alpha (8-10 Hz) and upper alpha (11-14 Hz). Bonferroni correction for multiple comparisons were employed to correct for these different frequency bands tested.

## Results

### Behavioral results

Perceptual judgments of the stimuli, presented randomly in the left or right visual hemifield, were bistable in both types of trials (two-flash fusion/FTT and apparent motion/AM). Specifically, two distinct flashes were reported on average on 40.2% (SD = 17.4%) of trials in the TFF condition and 46.3% (SD = 8.2%) of trials in the AM condition. The two trial types did not differ significantly in the rate of segregation/integration trials (t(_26_) = −1.6, p = .12). These results suggest that ISI values effectively caused the two stimuli to be integrated on about half of the trials.

### ERFs and cortical sources estimation

Cluster-based permutation tests allowed us to detect reliable differences between the ERFs evoked by segregation and integration in both the TFF and AM tasks. The complete set of sensors showing cluster-corrected significant differences for each comparison can be seen in 1B. In the TFF condition, ERFs started to differ as a function of subjective perception as early as 84 ms after the first flash onset (around 24 ms after both stimuli had offset) and continued till the end of the time period considered (700 ms) in a large group of sensors (minimum cluster-corrected p = .002; maximum cluster-corrected p = .033). In the AM condition, ERFs started to differ in a later time window, possibly because the second stimulus here appeared 120 ms after the first one; specifically, ERFs differed significantly starting from 390 ms (250 ms after both stimuli had been presented and removed) and continued till the end of the time period considered (700 ms; minimum/maximum cluster-corrected p = .049). As a general pattern emerging from the ERF analysis visible both in the TFF and the AM tasks, in all sensors where cluster-corrected differences emerged, two stimuli that were segregated elicited higher ERF amplitudes. Cortical sources estimation allowed us to identify ROIs that showed differential activity as a function of the type of percept (single/motion vs. double/alternation). They were considered as ROIs for pre-stimulus analyses (MVPA decoding and connectivity) only if their extension was equal or exceeded 10 cortical vertices.

The TFF and AM tasks elicited activities in two large and mostly non-overlapping cortical regions (Figure 2). Specifically, the TTF task showed different activity in visual areas including bilateral V1/V2 and the right MT area, in the left superior temporal area and bilaterally in frontal areas, i.e. the inferior and mid/superior frontal gyri. In contrast, the AM task showed different activity in the right MT area, in the right intraparietal sulcus IPS, in the right temporo-parietal junction (TPJ), in the right middle temporal gyrus, in the right insula and bilaterally in the superior frontal gyrus. The complete set of sources that for showed both tasks significant different activations are displayed and their anatomical labels are reported in Table 1 together with their MNI coordinates and cortical locations as derived from Automated Anatomical Labeling (AAL; Tzourio-Mazoyer et al. 2002).

**Figure 2:**
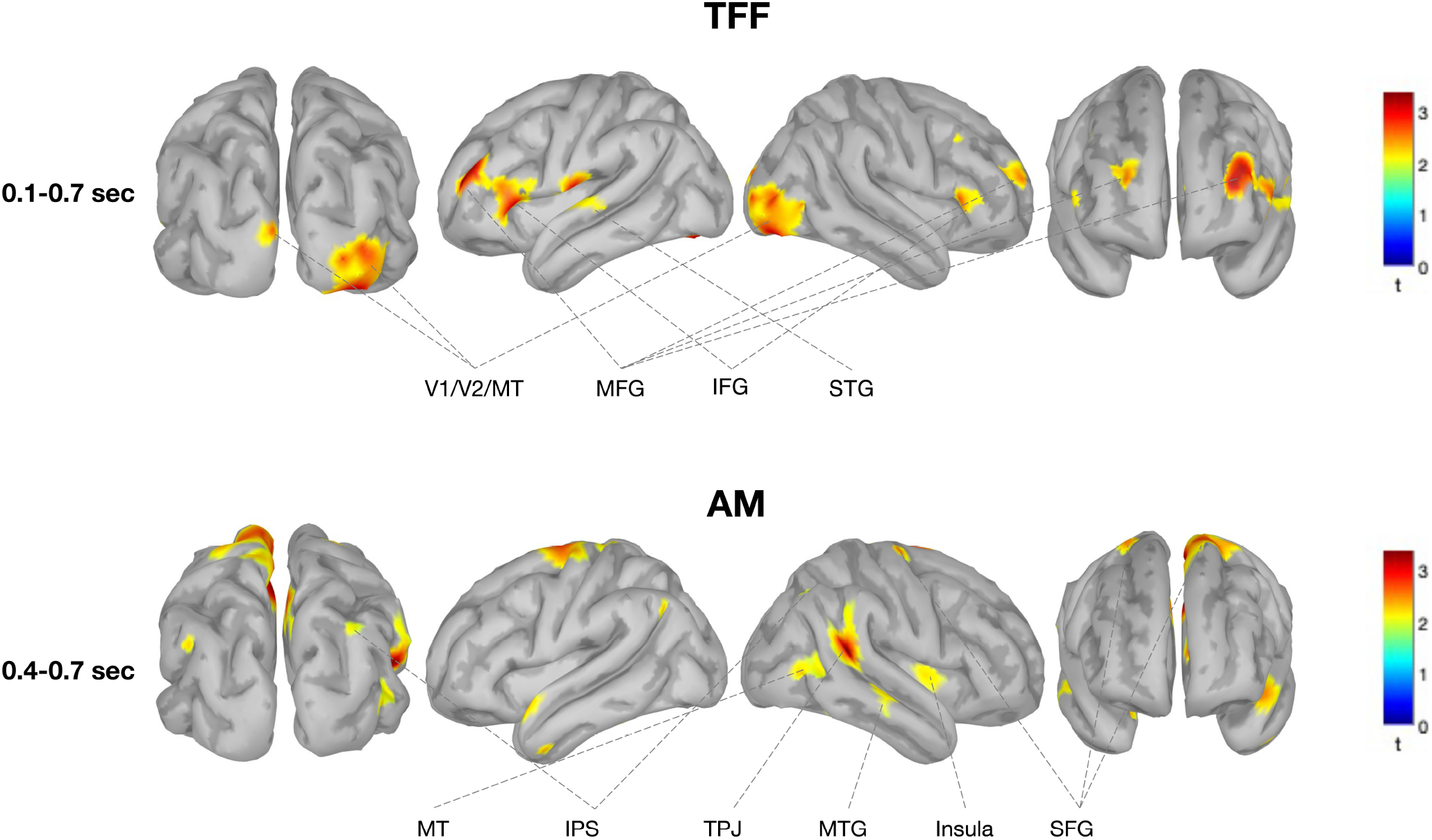
Event-related neural activity at the source level for the TFF and AM task. Cortical maps of activity (segregation–integration difference expressed as t-values) at the source level, averaged over the time windows where we found significant cluster-based permutation differences at the sensors level (from 100 to 700 ms after the first target onset for the TFF task and from 400 to 700 ms after the first target onset for the AM task). These areas that responded differently in the post-stimulus period have been considered ROIs for pre-stimulus MVPA and connectivity analyses if they passed the statistical threshold and also if extension was equal or above 10 cortical vertices. For details about the labelling and cortical locations of ROIs see Table 1.

### Decoding of perceptual outcome from prestimulus sources activity

Single-trials data from all ROIs were extracted and the relative time-frequency transformations were obtained to evaluate if prestimulus activity could be used to decode subjective perception, and if so, at which frequencies. The decoding accuracy (t-values) for the different perceptual outcomes obtained with the naive Bayes classifier searchlight performed on single-trial phase values is shown in Figures 3 and 4. Cluster-corrected permutation tests revealed that the time-frequency ranges in which subjective perception could be accurately decoded was different between the TFF and the AM task, and it was observed in different ROIs. Indeed, the highest decoding accuracy in predicting subjects’ perceptual outcome from the phase of prestimulus oscillation in the TFF task was found in the MT area of the right hemisphere, with frequencies spanning predominantly the theta and the alpha band (≈5-12 Hz) and around −400/-200 ms relative to the onset of the first stimulus (p=.048). On the contrary, decoding perceptual outcome in the AM condition was significant in the right IPS area for frequencies in theta band (≈4-7 Hz) at and around −700/-400 ms relative to the onset of the first flash (*p* = .026). Notably, these findings are perfectly in line in terms of time/frequency windows with previous EEG evidence in an independent participant sample (Ronconi et al., 2017), representing thus a replication of our previous findings.

**Figure 3:**
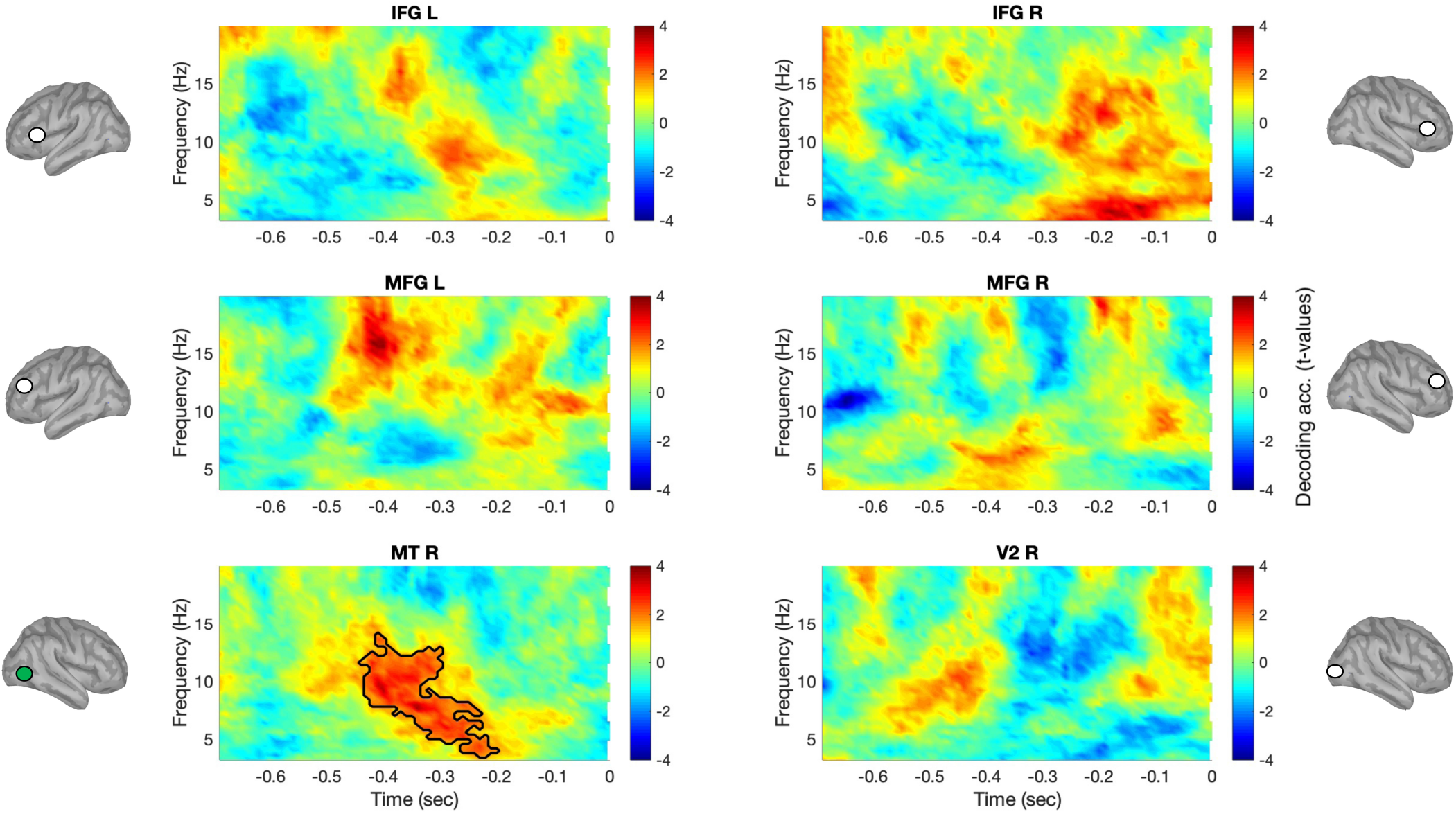
Single-trial prestimulus activity of the right MT area successfully decoded subjective perception in the TFF task. The prestimulus activity of source-level ROIs defined based on post-stimulus differences between perceptual outcome was the focus of a multivariate decoding analysis that aimed at evaluating whether ongoing phase at different oscillatory rhythms could predict subjective perception (i.e. temporal integration/segregation; single vs. double flash) in the TFF task (for both left and right visual hemifields). For each ROI, time-frequency plots show the group-level phase-decoding accuracy (difference against the chance level/50% decoding accuracy, expressed in t-values) obtained with a naive Bayes classifier searchlight. The outlined areas of the plots delimit the time/frequency points in which a significant difference was obtained with cluster-corrected permutation tests.

**Figure 4:**
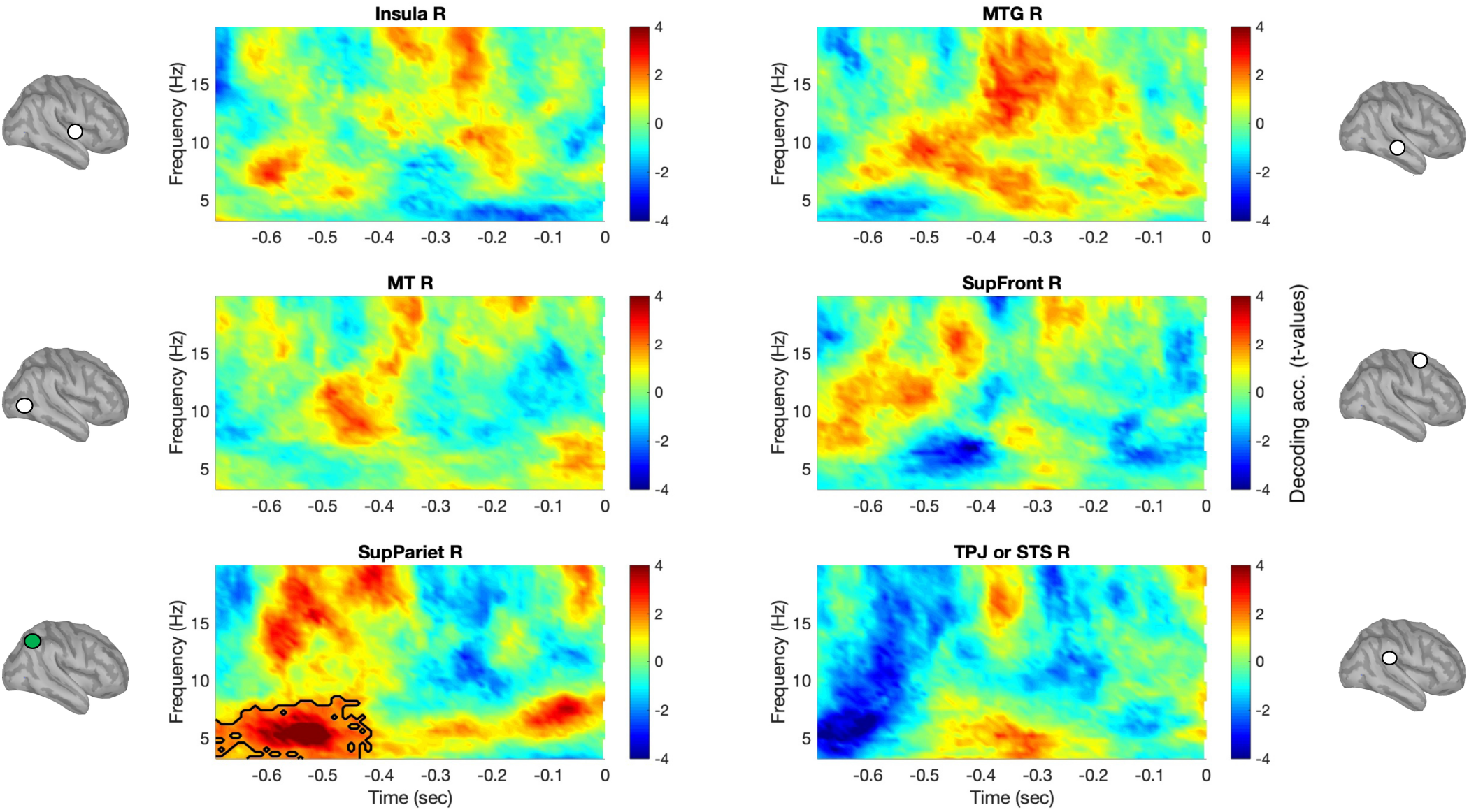
Single-trial prestimulus activity of the right superior parietal areas (including IPS) successfully decoded subjective perception in the AM task. The prestimulus activity of source-level ROIs defined based on post-stimulus differences between perceptual outcome was the focus of a multivariate decoding analysis that aimed at evaluating whether ongoing phase at different oscillatory rhythms could predict subjective perception (i.e. temporal integration/segregation; motion vs. alternation) in the AM task (for both left and right visual hemifields). For each ROI, time-frequency plots show the group-level phase-decoding accuracy (difference against the chance level/50% decoding accuracy, expressed in t-values) obtained with a naive Bayes classifier searchlight. The outlined areas of the plots delimit the time/frequency points in which a significant difference was obtained with cluster-corrected permutation tests.

### Prestimulus MEG connectivity is predictive of upcoming perceptual integration/segregation

Based on results of MVPA that revealed significant decoding performance from prestimulus activity of the right MT (for the TFF task) and of the right IPS (for the AM task) areas, we used these ROIs as hubs for pre-stimulus connectivity analyses within the extended network that showed differential activation as a function of integration/segregation of visual stimuli. For the TFF task (Figure 5), we found that perceptual segregation (i.e. perception of two distinct flashes) was preceded by a significant increase of prestimulus connectivity in the upper alpha band (11-14 Hz) between the areas MT and V2 of the right hemisphere (p = 0.0384; one-tailed, Bonferroni corrected); similarly, a tendency for a significant increment in pre-stimulus connectivity was found in the theta band (4-7 Hz) between the areas MT and IFG of the right hemisphere (p = 0.0505; one-tailed, Bonferroni corrected). No other pairwise differences in connectivity to/from the area MT were found to be significant (all p > .289).

**Figure 5:**
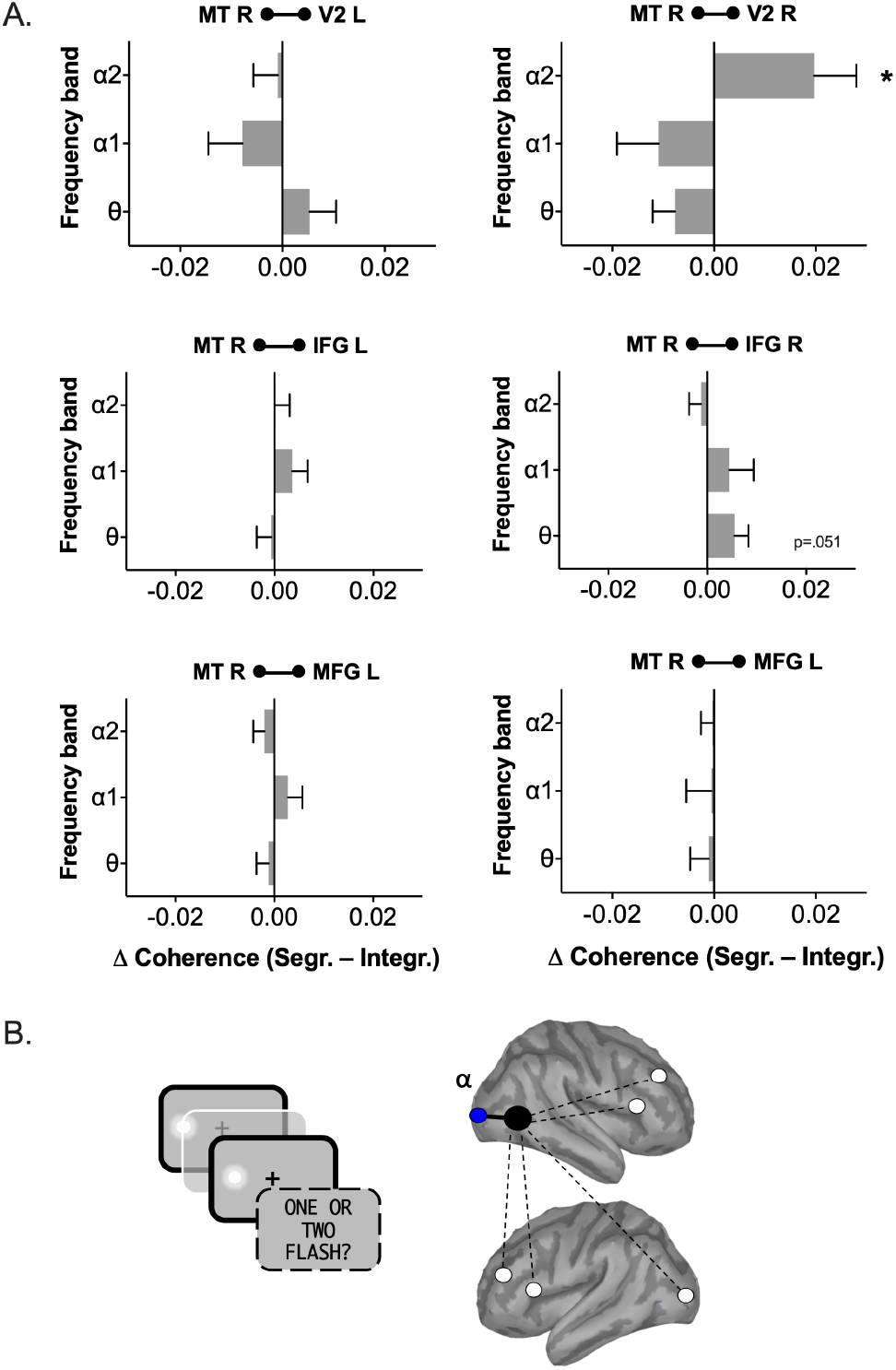
Connectivity results for the TFF task. (A) Following the results obtained in the MVPA analysis, we used the right MT area as the seed for testing functional connectivity (based on phase coherence) between this area and the other ROIs differentially activated in the TFF task. Results showed a significant increment of connectivity before the onset of the targets between right MT and early visual areas (i.e. V2) in segregation trials, as compared to integration trials, in the upper alpha band (11-14 Hz). A trend toward a significant increment in connectivity in segregation trials was found also in slower frequencies in the theta band between right MT and right IFG areas. (B) Schematic representation of connectivity results obtained, where bold lines mean significant connectivity variation between pairs of relevant ROIs, while dotted lines are displayed where no significant connectivity differences were obtained.

For the AM task (Figure 6), we found that perceptual segregation (i.e. perception of two distinct flashes) was preceded by a significant increase of prestimulus connectivity in the theta band (4-7 Hz) between the areas IPS/Superior Parietal and MT of the right hemisphere (p = .045; one-tailed, Bonferroni corrected). No other pairwise differences in connectivity to/from the right IPS/Superior Parietal were found to be significant (all p > .087).

**Figure 6:**
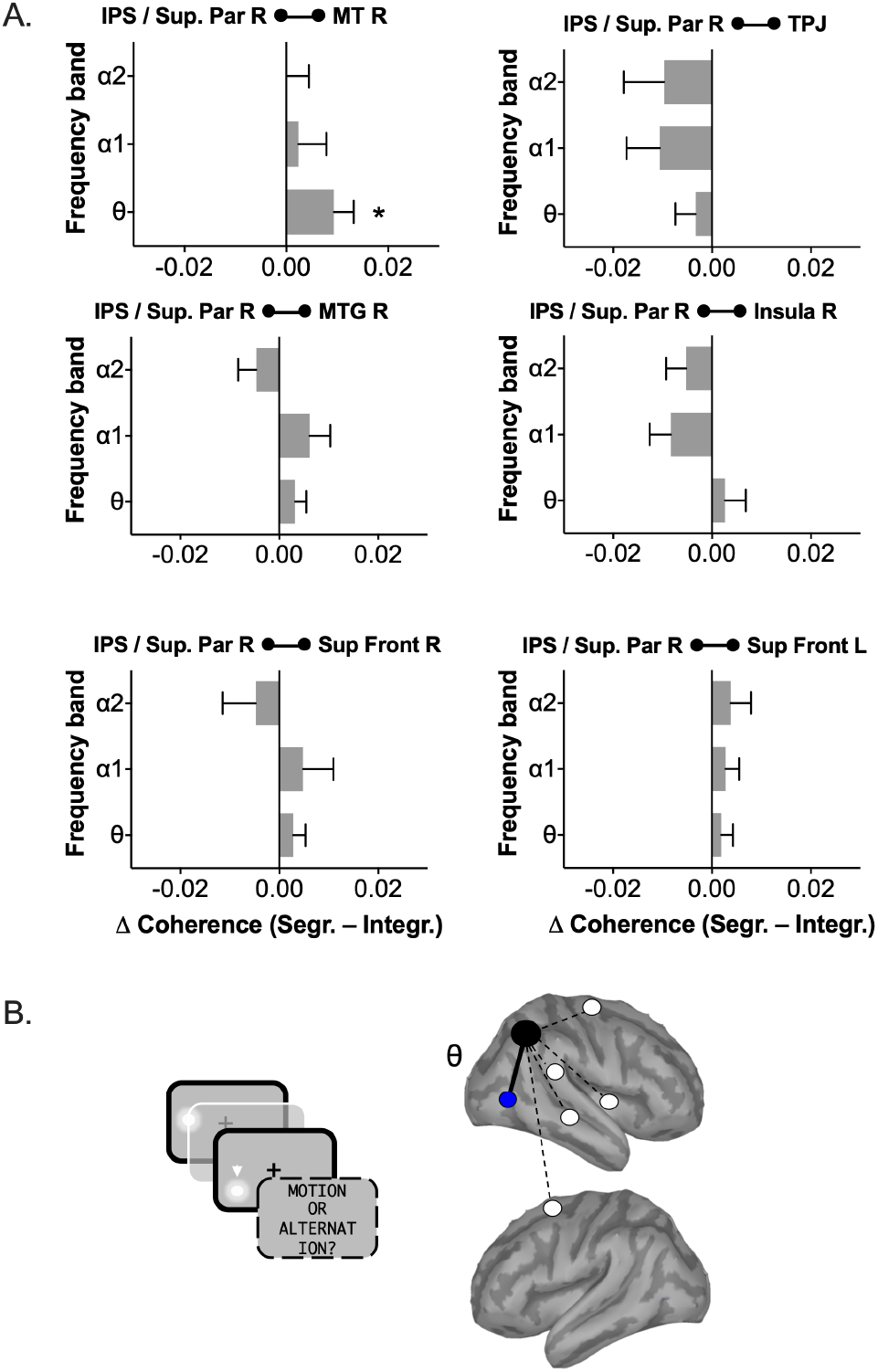
Connectivity results for the AM task. (A) Following the results obtained in the MVPA analysis, we used the right IPS/Superior Parietal area as the seed for testing functional connectivity (based on phase coherence) between this area and the other ROIs differentially activated in the AM task. Results showed a significant increment of connectivity before the onset of the targets between the right IPS/Superior Parietal and the right MT are in segregation trials, as compared to integration trials, in the theta band (4-7 Hz). (B) Schematic representation of connectivity results obtained, where bold lines mean significant connectivity variation between pairs of relevant ROIs, while dotted lines are displayed where no significant connectivity differences were obtained.

## Discussion

Starting from the idea that one cycle of low-frequency neural oscillation represents the elementary unit for sampling sensory information in different domains (Poppel, 1997; Van Wassenhove, 2016; VanRullen, 2016), in the present study we used multivariate decoding of MEG data to shed light on the neural networks underlying the fundamental ability of the human visual system to integrate and segregate visual input. Our findings clearly point to two different functional networks underlying two aspects of visual temporal processing. The first network, involved in rapid temporal segregation of stimuli separated by just a few tens of milliseconds, was associated with early visual processing areas and visual area V5/MT. Indeed, V5/MT is sensitive to stimuli presented at high temporal frequency and has been previously associated with temporal perception (Bueti et al., 2008a). Here, we showed that the phase of alpha oscillations localized to this area predicted integration versus segregation in the two-flash fusion task. Moreover, V5/MT also showed increased functional connectivity with early visual areas (V2) in the upper alpha-band when participant segregated the two stimuli.

In contrast, for a longer temporal scale and with visual information displayed in different spatial locations, higher-order cortical areas in the parietal lobe (i.e. IPS/Superior parietal cortex) were identified as the source of phase decoding in the theta band. This area showed also increased theta-band connectivity with the area V5/MT when participant segregated the two stimuli as opposed to perceiving a single object in (apparent) motion. These findings build on work showing a prominent theta band rhythm in visual processing areas (Spyropoulos et al., 2018) as well as in parietal cortex (Raghavachari et al., 2006), and suggest the active integration and segregation of sensory stimuli, at least in the visual modality, relies on a phase-dependent temporal coding at low-mid frequency oscillations. Thanks to principles of phase-amplitude coupling previously demonstrated for both alpha and theta oscillations (Jensen et al., 2014; Lisman and Jensen, 2013; Koster et al., 2019), low-mid frequency oscillations would then modulate gamma-band activity in order to organize simple perceptual representations in time, limiting the number of representations that can be processed in each oscillatory cycle depending on ‘hardware’ limits, i.e. the basic temporal resolution of our visual system, and also on whether they involve tracking of events in a single or different spatial location. The co-existence of these different rhythms could theoretically account also for integration of stimuli of higher complexity than the ones employed in the present study, such as words, objects or faces (Drewes et al., 2015; Wang and Luo, 2017), that would require a more complex brain network of visual regions to be tracked in their spatiotemporal dynamics (Baldauf and Desimone, 2014; de Vries and Baldauf, 2019).

Our results are among the first to elucidate cortical origins of alpha and theta activity in the context of visual temporal parsing, building on previous sensor-level EEG findings (Ronconi et al., 2017). In fact, in the present study we replicate, in a new set of participants and with different neuroimaging tools, our previous EEG finding showing that the perceptual interpretation (integration vs. segregation) depended on the phase of ongoing/prestimulus oscillations at different frequency bands (Ronconi et al., 2017). Not only were the frequencies showing maximum decoding accuracy for the tasks closely matching between the present MEG and the previous EEG data, but there was a matching topography of the maximum decoding accuracy with the right posterior channels found already previously compatible with results obtained here at the cortical sources level. The current findings replicate and substantially extend those findings to also uncover the network connectivity that may underlie these two sampling frequencies. Our results are in line with previous theoretical proposal claiming that timing does not involve a single, centralized clock for the visual system, but that visual timing is dependent on the pattern of activity within distributed networks (Burr and Morrone, 2006). In addition, our findings agree with TMS evidence causally linking V5/MT to timing processes in the visual domain that have used temporal discrimination tasks (Bueti et al., 2008b,a; Salvioni et al., 2013; Mioni et al., 2020).

Moreover, the evidence reported here are in line with other studies showing dissociations between MT and parietal lobe for spatiotemporal resolution of perception and motion extrapolation (Battelli et al., 2003, 2001) and with theoretical models proposing the existence of a “when” pathway in the human visual system involving V5/MT and the parietal lobe of the right hemisphere (Battelli et al., 2007). Interestingly, patients with right parietal damage do not have impairment in low-level temporal processing as measured by flicker detection thresholds (Battelli et al., 2003). In contrast, parietal patients have shown a bilateral deficit in apparent motion perception (Battelli et al., 2001), while deficits in other attentional tasks, such as multiple-object tracking, were present only in the hemifield contralateral to the parietal lesion. Such dependence of apparent motion perception on right hemisphere areas – irrespective of stimuli presentation hemifield – closely matches our data showing the involvement of a network of right hemispheric regions. Together, these results suggest that the parietal cortex of the right hemisphere may serve as a main control hub for theta-driven spatiotemporal integration in visual perception.

The definition of these different networks at the source-level might stimulate future transcranial electrical stimulation (tES) studies, in an attempt to bring causal evidence about the role of oscillatory activity within these networks in determining spatio-temporal aspects of perception. In particular, transcranial alternating current stimulation (tACS) could be of central relevance due to its ability to modulate brain oscillations in a frequency-dependent manner, with initial evidence linking tACS at specific frequencies with some basics binding processes in the visual system (for a review see Ghiani et al. 2021). Using the two-flash fusion tasks, (Battaglini et al., 2020) showed recently that participants tend to integrate two subsequent flashes more often (i.e., they tend to report just one flash) when 10 Hz tACS (i.e., alpha tACS) was applied over V5/MT of the right hemisphere and surrounding extrastriate visual regions. On the contrary, 18Hz tACS (i.e., tACS within the beta band) and sham had no significant effect on the reported number of flashes. These findings provide initial confirmation to the idea of a causal link between alpha activity in V5/MT and extra-striate visual regions and temporal integration/segregation for short temporal scales.

The importance of the current replication and extension of previous results using this same (Ronconi and Melcher, 2017; Ronconi et al., 2017) or similar paradigms (Varela et al., 1981; Mathewson et al., 2009; Wutz et al., 2014, 2016; Milton and Pleydell-Pearce, 2016) is heightened by multiple null findings in recent studies investigating the impact of ongoing alpha oscillations on perception. In fact, several recent works show no effect of alpha phase in stimuli detection (Ruzzoli et al., 2019), visual awareness and accuracy (Benwell et al., 2017, 2021) or reaction times (Vigué-Guix et al., 2020). In another study using both flashes and sounds, Buergers and Noppeney (2022) found no effect of alpha frequency (both as an individual trait and as a varying state) on visual integration, posing an important challenge to the claim that ongoing alpha oscillations impact the temporal precision of visual perception.

Given the set of null findings just cited, the relevance of the current work is twofold: firstly, it reinforces the idea that the phase of ongoing alpha band oscillations shapes our conscious perception, and that this contribution is critical in the integration and segregation of visual stimuli. Secondly, contrarily to Buergers and Noppeney (2022), we provide evidence of the role of ongoing alpha oscillations in pacing visual perception, by demonstrating a pattern of connectivity between V2 and MT critically specific to the upper alpha band and to the segregation of visual stimuli. Reasons of this discrepancy might have to be searched into the different sources of alpha oscillations (Womelsdorf et al., 2014), characterizing either topdown cortical influences (Van Kerkoerle et al., 2014; Halgren et al., 2019) or thalamo-cortical communications (Bollimunta et al., 2011; Hughes et al., 2011).

To summarize, the current results demonstrate the existence of two networks for visual temporal integration: an alpha frequency network involving relatively early visual processing areas that determines the temporal resolution of rapid visual perception, and a slower, theta frequency network involving parietal regions in the interpretation of more complex spatiotemporal events. The different sampling frequencies, alpha and theta, may reflect different network activity and connectivity patterns in right-lateralized cortical regions that form the human dorsal “where” and “when” systems.

## References

Baldauf, D. and Desimone, R. (2014). Neural mechanisms of object-based attention. Science, 344(6182):424–427.

Battaglini, L., Ghiani, A., Casco, C., and Ronconi, L. (2020). Parietal tacs at beta frequency improves vision in a crowding regime. Neuroimage, 208:116451.

Battelli, L., Cavanagh, P., Intriligator, J., Tramo, M. J., Hénaff, M.-A., Michèl, F., and Barton, J. J. (2001). Unilateral right parietal damage leads to bilateral deficit for high-level motion. Neuron, 32(6):985–995.

Battelli, L., Cavanagh, P., and Thornton, I. M. (2003). Perception of biological motion in parietal patients. Neuropsychologia, 41(13):1808–1816.

Battelli, L., Pascual-Leone, A., and Cavanagh, P. (2007). The ‘when’pathway of the right parietal lobe. Trends in cognitive sciences, 11(5):204–210.

Benwell, C. S., Coldea, A., Harvey, M., and Thut, G. (2021). Low pre-stimulus EEG alpha power amplifies visual awareness but not visual sensitivity. European Journal of Neuroscience.

Benwell, C. S., Tagliabue, C. F., Veniero, D., Cecere, R., Savazzi, S., and Thut, G. (2017). Prestimulus EEG power predicts conscious awareness but not objective visual performance. Eneuro, 4(6).

Berens, P. (2009). Circstat: a matlab toolbox for circular statistics. Journal of statistical software, 31:1–21.

Bishop, G. H. (1932). Cyclic changes in excitability of the optic pathway of the rabbit. American Journal of Physiology, 103(1):213–224.

Bollimunta, A., Mo, J., Schroeder, C. E., and Ding, M. (2011). Neuronal mechanisms and attentional modulation of corticothalamic alpha oscillations. Journal of Neuroscience, 31(13):4935–4943.

Buergers, S. and Noppeney, U. (2022). The role of alpha oscillations in temporal binding within and across the senses. Nature Human Behaviour, pages 1–11.

Bueti, D., Bahrami, B., and Walsh, V. (2008a). Sensory and association cortex in time perception. Journal of Cognitive Neuroscience, 20(6):1054–1062.

Bueti, D., van Dongen, E. V., and Walsh, V. (2008b). The role of superior temporal cortex in auditory timing. PLoS One, 3(6):e2481.

Burr, D. and Morrone, C. (2006). Time perception: space–time in the brain. Current Biology, 16(5):R171–R173.

Cecere, R., Rees, G., and Romei, V. (2015). Individual differences in alpha frequency drive crossmodal illusory perception. Current Biology, 25(2):231–235.

de Vries, E. and Baldauf, D. (2019). Attentional weighting in the face processing network: a magnetic response image-guided magnetoencephalography study using multiple cyclic entrainments. Journal of cognitive neuroscience, 31(10):1573–1588.

Drewes, J., Zhu, W., Wutz, A., and Melcher, D. (2015). Dense sampling reveals behavioral oscillations in rapid visual categorization. Scientific Reports, 5(1):1–9.

Fischl, B. (2012). Freesurfer. Neuroimage, 62(2):774–781.

Ghiani, A., Maniglia, M., Battaglini, L., Melcher, D., and Ronconi, L. (2021). Binding mechanisms in visual perception and their link with neural oscillations: A review of evidence from tacs. Frontiers in Psychology, 12:779.

Gulbinaite, R., İlhan, B., and VanRullen, R. (2017). The triple-flash illusion reveals a driving role of alpha-band reverberations in visual perception. Journal of Neuroscience, 37(30):7219–7230.

Haegens, S., Nácher, V., Luna, R., Romo, R., and Jensen, O. (2011). *α*-oscillations in the monkey sensorimotor network influence discrimination performance by rhythmical inhibition of neuronal spiking. Proceedings of the National Academy of Sciences, 108(48):19377–19382.

Halgren, M., Ulbert, I., Bastuji, H., Fabó, D., Erőss, L., Rey, M., Devinsky, O., Doyle, W. K., Mak-McCully, R., Halgren, E., et al. (2019). The generation and propagation of the human alpha rhythm. Proceedings of the National Academy of Sciences, 116(47):23772–23782.

Hämäläinen, M. S. and Ilmoniemi, R. J. (1994). Interpreting magnetic fields of the brain: minimum norm estimates. Medical & biological engineering & computing, 32(1):35–42.

Harter, M. R. (1967). Excitability cycles and cortical scanning: a review of two hypotheses of central intermittency in perception. Psychological bulletin, 68(1):47.

Huang, M., Mosher, J. C., and Leahy, R. (1999). A sensor-weighted overlapping-sphere head model and exhaustive head model comparison for meg. Physics in Medicine & Biology, 44(2):423.

Hughes, S., Lőrincz, M., Turmaine, M., and Crunelli, V. (2011). Thalamic gap junctions control local neuronal synchrony and influence macroscopic oscillation amplitude during EEG alpha rhythms. Frontiers in Psychology, 2:193.

Jensen, O., Gips, B., Bergmann, T. O., and Bonnefond, M. (2014). Temporal coding organized by coupled alpha and gamma oscillations prioritize visual processing. Trends in neurosciences, 37(7):357–369.

Köster, M., Martens, U., and Gruber, T. (2019). Memory entrainment by visually evoked theta-gamma coupling. NeuroImage, 188:181–187.

Lansing, R. W. (1957). Relation of brain and tremor rhythms to visual reaction time. Electroencephalography and clinical neurophysiology, 9(3):497–504.

Lisman, J. E. and Jensen, O. (2013). The theta-gamma neural code. Neuron, 77(6):1002–1016.

Mathewson, K. E., Gratton, G., Fabiani, M., Beck, D. M., and Ro, T. (2009). To see or not to see: prestimulus *α* phase predicts visual awareness. Journal of Neuroscience, 29(9):2725–2732.

Milton, A. and Pleydell-Pearce, C. W. (2016). The phase of pre-stimulus alpha oscillations influences the visual perception of stimulus timing. Neuroimage, 133:53–61.

Mioni, G., Grondin, S., Bardi, L., and Stablum, F. (2020). Understanding time perception through non-invasive brain stimulation techniques: A review of studies. Behavioural brain research, 377:112232.

Morillon, B. and Schroeder, C. E. (2015). Neuronal oscillations as a mechanistic substrate of auditory temporal prediction. Annals of the New York Academy of Sciences, 1337(1):26–31.

Nolte, G. (2003). The magnetic lead field theorem in the quasistatic approximation and its use for magnetoencephalography forward calculation in realistic volume conductors. Physics in Medicine & Biology, 48(22):3637.

Oostenveld, R., Fries, P., Maris, E., and Schoffelen, J.-M. (2011). Fieldtrip: open source software for advanced analysis of meg, EEG, and invasive electrophysiological data. Computational intelligence and neuroscience, 2011.

Oosterhof, N. N., Connolly, A. C., and Haxby, J. V. (2016). Cosmomvpa: multi-modal multivariate pattern analysis of neuroimaging data in matlab/gnu octave. Frontiers in neuroinformatics, 10:27.

Pöppel, E. (1997). A hierarchical model of temporal perception. Trends in cognitive sciences, 1(2):56–61.

Raghavachari, S., Lisman, J. E., Tully, M., Madsen, J. R., Bromfield, E., and Kahana, M. J. (2006). Theta oscillations in human cortex during a working-memory task: evidence for local generators. Journal of neurophysiology, 95(3):1630–1638.

Rassi, E., Wutz, A., Müller-Voggel, N., and Weisz, N. (2019). Prestimulus feedback connectivity biases the content of visual experiences. Proceedings of the National Academy of Sciences, 116(32):16056–16061.

Ronconi, L., Busch, N. A., and Melcher, D. (2018). Alpha-band sensory entrainment alters the duration of temporal windows in visual perception. Scientific reports, 8(1):1–10.

Ronconi, L. and Melcher, D. (2017). The role of oscillatory phase in determining the temporal organization of perception: evidence from sensory entrainment. Journal of Neuroscience, 37(44):10636–10644.

Ronconi, L., Oosterhof, N. N., Bonmassar, C., and Melcher, D. (2017). Multiple oscillatory rhythms determine the temporal organization of perception. Proceedings of the National Academy of Sciences, 114(51):13435–13440.

Ruzzoli, M., Torralba, M., Fernández, L. M., and Soto-Faraco, S. (2019). The relevance of alpha phase in human perception. Cortex, 120:249–268.

Salvioni, P., Murray, M. M., Kalmbach, L., and Bueti, D. (2013). How the visual brain encodes and keeps track of time. Journal of Neuroscience, 33(30):12423–12429.

Samaha, J. and Postle, B. R. (2015). The speed of alpha-band oscillations predicts the temporal resolution of visual perception. Current Biology, 25(22):2985–2990.

Schoffelen, J.-M. and Gross, J. (2009). Source connectivity analysis with meg and EEG. Human brain mapping, 30(6):1857–1865.

Spyropoulos, G., Bosman, C. A., and Fries, P. (2018). A theta rhythm in macaque visual cortex and its attentional modulation. Proceedings of the National Academy of Sciences, 115(24):E5614–E5623.

Tadel, F., Baillet, S., Mosher, J. C., Pantazis, D., and Leahy, R. M. (2011). Brainstorm: a user-friendly application for meg/EEG analysis. Computational intelligence and neuroscience, 2011.

Tadel, F., Bock, E., Niso, G., Mosher, J. C., Cousineau, M., Pantazis, D., Leahy, R. M., and Baillet, S. (2019). Meg/EEG group analysis with brainstorm. Frontiers in neuroscience, page 76.

Taulu, S. and Kajola, M. (2005). Presentation of electromagnetic multichannel data: the signal space separation method. Journal of Applied Physics, 97(12):124905.

Taulu, S., Simola, J., and Kajola, M. (2005). Applications of the signal space separation method. IEEE transactions on signal processing, 53(9):3359–3372.

Tzourio-Mazoyer, N., Landeau, B., Papathanassiou, D., Crivello, F., Etard, O., Delcroix, N., Mazoyer, B., and Joliot, M. (2002). Automated anatomical labeling of activations in spm using a macroscopic anatomical parcellation of the mni mri single-subject brain. Neuroimage, 15(1):273–289.

Van Kerkoerle, T., Self, M. W., Dagnino, B., Gariel-Mathis, M.-A., Poort, J., Van Der Togt, C., and Roelfsema, P. R. (2014). Alpha and gamma oscillations characterize feedback and feedforward processing in monkey visual cortex. Proceedings of the National Academy of Sciences, 111(40):14332–14341.

Van Wassenhove, V. (2016). Temporal cognition and neural oscillations. Current Opinion in Behavioral Sciences, 8:124–130.

VanRullen, R. (2016). Perceptual cycles. Trends in cognitive sciences, 20(10):723–735.

Varela, F. J., Toro, A., John, E. R., and Schwartz, E. L. (1981). Perceptual framing and cortical alpha rhythm. Neuropsychologia, 19(5):675–686.

Vigué-Guix, I., Morís Fernández, L., Torralba Cuello, M., Ruzzoli, M., and Soto-Faraco, S. (2020). Can the occipital alpha-phase speed up visual detection through a realtime EEG-based brain–computer interface (bci)? European Journal of Neuroscience.

Wang, Y. and Luo, H. (2017). Behavioral oscillation in face priming: Prediction about face identity is updated at a theta-band rhythm. Progress in Brain Research, 236:211–224.

White, P. A. (2018). Is conscious perception a series of discrete temporal frames? Consciousness and cognition, 60:98–126.

Womelsdorf, T., Valiante, T. A., Sahin, N. T., Miller, K. J., and Tiesinga, P. (2014). Dynamic circuit motifs underlying rhythmic gain control, gating and integration. Nature neuroscience, 17(8):1031–1039.

Wutz, A., Melcher, D., and Samaha, J. (2018). Frequency modulation of neural oscillations according to visual task demands. Proceedings of the National Academy of Sciences, 115(6):1346–1351.

Wutz, A., Muschter, E., van Koningsbruggen, M. G., Weisz, N., and Melcher, D. (2016). Temporal integration windows in neural processing and perception aligned to saccadic eye movements. Current biology, 26(13):1659–1668.

Wutz, A., Weisz, N., Braun, C., and Melcher, D. (2014). Temporal windows in visual processing:”prestimulus brain state” and “poststimulus phase reset” segregate visual transients on different temporal scales. Journal of Neuroscience, 34(4):1554–1565.

